# Versatile and robust genome editing with *Streptococcus thermophilus* CRISPR1-Cas9

**DOI:** 10.1101/321208

**Authors:** Daniel Agudelo, Sophie Carter, Minja Velimirovic, Alexis Duringer, Sébastien Levesque, Jean-François Rivest, Jeremy Loehr, Mathilde Mouchiroud, Denis Cyr, Paula J Waters, Mathieu Laplante, Sylvain Moineau, Adeline Goulet, Yannick Doyon

## Abstract

Targeting definite genomic locations using CRISPR-Cas systems requires a set of enzymes with unique protospacer adjacent motif (PAM) compatibilities. To expand this repertoire, we engineered nucleases, cytosine base editors, and adenine base editors from the archetypal *Streptococcus thermophilus* CRISPR1-Cas9 (St1Cas9) system. We found that St1Cas9 strain variants enable targeting to five distinct A-rich PAMs and provide structural basis for their specificities. The small size of this ortholog enables expression of the holoenzyme from a single adeno-associated viral vector for *in vivo* editing applications. Delivery of St1Cas9 to the neonatal liver efficiently rewired metabolic pathways, leading to phenotypic rescue in a mouse model of hereditary tyrosinemia. These robust enzymes expand and complement current editing platforms available for tailoring mammalian genomes.

## INTRODUCTION

Clustered regularly interspaced short palindromic repeats (CRISPR) and CRISPR-associated (Cas) proteins form a prokaryotic adaptive immune system and some of its components have been harnessed for robust genome editing (Komor et al., 2017). Type II-based editing tools rely on a large multidomain endonuclease, Cas9, guided to its DNA target by an engineered single-guide RNA (sgRNA) chimera (Jinek et al., 2012) (See (Koonin et al., 2017; Makarova et al., 2018; Shmakov et al., 2017) for a classification of CRISPR-Cas systems). The Cas9-sgRNA binary complex finds its target through recognition of a short sequence called the protospacer adjacent motif (PAM) and the subsequent base pairing between the guide RNA and DNA leads to a double-strand break (DSB) (Hille et al., 2018; Komor et al., 2017). While *Streptococcus pyogenes* (SpCas9) remains the most widely used Cas9 ortholog for genome engineering, the diversity of naturally occurring RNA-guided nucleases is astonishing (Shmakov et al., 2017). Hence, Cas9 enzymes from different microbial species can contribute to the expansion of the CRISPR toolset by increasing targeting density, improving activity and specificity as well as easing delivery (Esvelt et al., 2013; Komor et al., 2017).

In principle, engineering complementary CRISPR-Cas systems from distinct bacterial species should be relatively straightforward, as they have been minimized to only two components. However, many such enzymes were found inactive in human cells despite being accurately reprogrammed for DNA binding and cleavage *in vitro* (Chen et al., 2017; Ran et al., 2015; Zetsche et al., 2015). Nevertheless, a striking example of the value of alternative Cas9 enzymes is the implementation of the type II-A Cas9 from *Staphylococcus aureus* (SaCas9) for *in vivo* editing using a single recombinant adeno-associated virus (AAV) vector (Ran et al., 2015). More recently, *Campylobacter jejuni* and *Neisseria meningitidis* Cas9s from the type II-C (Mir et al., 2018) CRISPR-Cas systems have been added to this repertoire (Edraki et al., 2019; Ibraheim et al., 2018; Kim et al., 2017). *In vivo* editing offers the possibility to generate phenotypes in animal models in order to better recapitulate the interactions between cell types and organs. In addition, it can be envisioned as a novel class of human therapeutics that enables precise molecular correction of genetic defects underlying diseases. Therefore, further development of robust and wide-ranging CRISPR-based technologies for *in vivo* editing may help to decipher disease mechanisms and offer novel therapeutic options (Lau and Suh, 2017; Schneller et al., 2017).

Here we revisited the properties of *Streptococcus thermophilus* type II-A CRISPR1-Cas9 system, a model nuclease of paramount importance to the entire CRISPR field (Barrangou and Horvath, 2017; Hille et al., 2018), and engineered a potent RNA-guided nuclease for both *in vitro* and *in vivo* applications. The distinctive functional PAM sequences of St1Cas9 variants increase the targeting flexibility and combinatorial potential of CRISPR-based nucleases and base editors. *In vivo* delivery using an all-in-one AAV led to efficient gene inactivation in the liver of neonatal mice, efficiently perturbing metabolic pathways. Taken together, these results highlight the benefits of engineering Cas9 orthologs, such as St1Cas9, to increase the versatility of CRISPR-based genome editing technologies.

## RESULTS

### Robust and potent DNA cleavage by St1Cas9 in human cells

While characterizing the interplay between St1Cas9 and anti-CRISPR proteins isolated from phages infecting *S. thermophilus* (Hynes et al., 2018) we noticed the substantial levels of editing achieved in human cells, an observation contrasting with previous reports (Chari et al., 2015; Ran et al., 2015). We thus revisited its properties and attempted to optimize its activity. First, we flanked the human codon-optimized ORF (Kleinstiver et al., 2015b) with nuclear localization signals (NLS) (**Figure 1A)**. Second, we customized the sgRNA sequence to maximize nuclease activity and tested our constructs at three endogenous loci (**Figures 1A-1B** and **S1A-1F**). The best performing sgRNA architecture (v1) was engineered by truncating the repeat:anti-repeat region (Briner et al., 2014) and substituting a wobble base pair present in the lower stem for a canonical Watson-Crick base pair (**Figure 1B**). These modifications also markedly improved transcriptional activation using dSt1Cas9-VPR (Chavez et al., 2015) (**Figure S2**). This analysis revealed that high gene disruption rates could be obtained under standard conditions using St1Cas9 in human cells.

**Figure 1.**
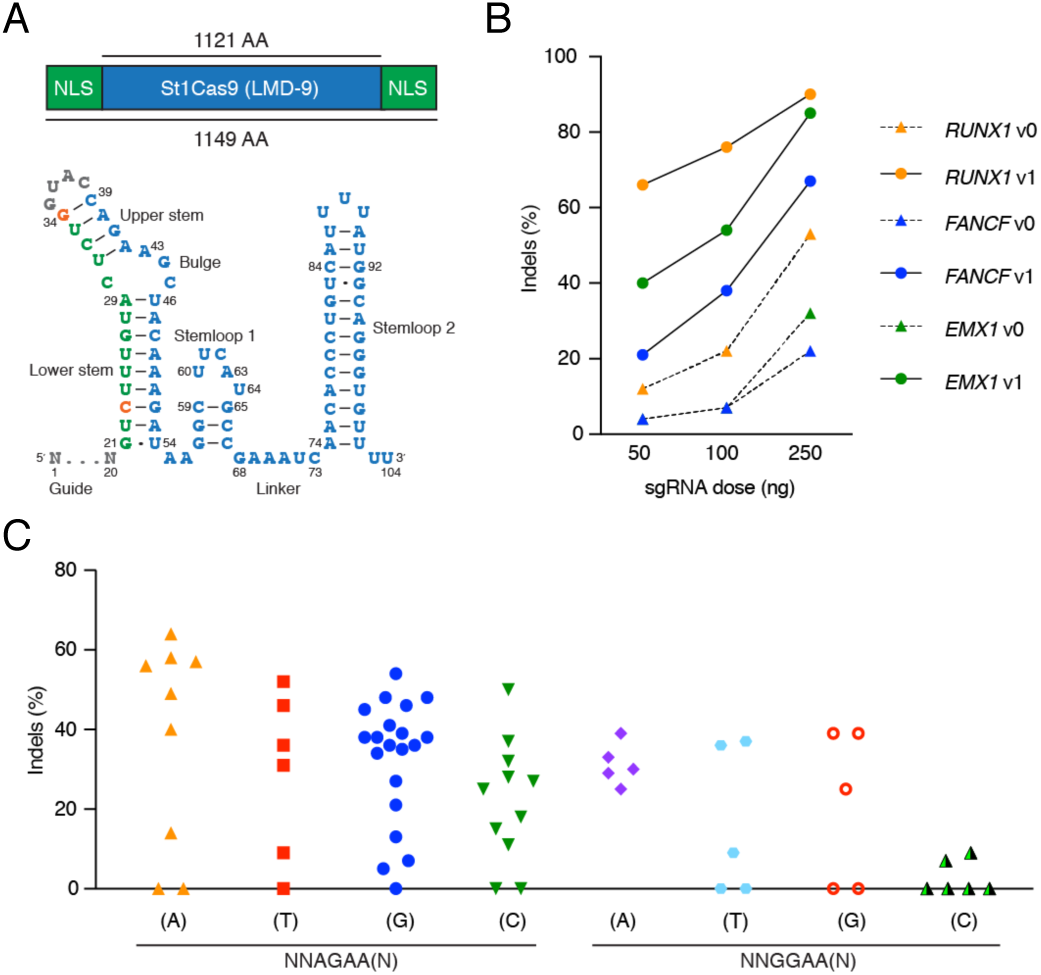
Functional PAM sequences for robust and potent DNA cleavage by St1Cas9 LMD-9 in mammalian cells. (A) Schematic representations of St1Cas9 LMD-9 flanked by nuclear localization signals (NLS) and its engineered sgRNA (v1). Nucleotide sequence and functional modules are depicted; crRNA (green), loop (grey), tracrRNA (blue), mutated nucleotides (orange). (B) K562 cells stably expressing St1Cas9 were transfected with indicated sgRNA expression vectors at increasing doses and TIDE assays were performed 3 days later to determine the frequency of indels. An expression vector encoding EGFP (-) was used as a negative control. The experiment was performed twice and yielded equivalent results. (C) Screening for guides targeting St1Cas9 LMD-9 to various PAMs was done by transient transfections in K562 and Neuro-2a cells using single vector constructs driving the expression of St1Cas9 and its sgRNA. Surveyor assays were performed 3 days later to determine the frequency of indels. An expression vector encoding EGFP (-) was used as a negative control. (See also **Figures S1** and **S2S**).

### Functional PAM sequences for St1Cas9 LMD-9 in mammalian cells

Cas9 orthologs rely on different PAMs as the first step in target recognition and the consensus PAM for St1Cas9 (LMD-9 and DGCC7710 *S. thermophilus* strains that differ by only 2 aa within their N-terminus) was originally defined as N_1_N_2_A_3_G_4_A_5_A_6_W_7_ (where W is A or T) (Deveau et al., 2008; Horvath et al., 2008). However, sequences closely related to the consensus can be functional in test tubes or when transplanted in *E. coli* for St1Cas9 LMD-9, the strain variant first engineered for genome editing (Cong et al., 2013; Esvelt et al., 2013; Kleinstiver et al., 2015b; Leenay et al., 2016). We thus explored its PAM preference by targeting endogenous loci in human and mouse cells. This analysis revealed that St1Cas9 LMD-9 functions efficiently at both NNAGAA and NNGGAA PAMs (**Figure 1C**). While a C is tolerated at position 7, there is a trend for these guides to be less efficacious. This bias was also observed in bacterial cells (Leenay et al., 2016). Thus, the functional core PAM sequence is constituted of four specific base pairs and defined as NNRGAA (where R is A or G). The optimal PAM sequence to regularly achieve high levels of editing is NNRGAAD (where D is A or G or T). The length of the nonconserved PAM linker (NN) has also been shown to be flexible and an extension from 2 to 3 bases can be tolerated in bacterial cells (Briner et al., 2014; Chen et al., 2014), but we failed to reproduce this observation in human cells suggesting a higher stringency of the system (**Figure S1G**). We also explored the impact of varying guide length on activity and observed no obvious correlation, confirming previous observations (Kleinstiver et al., 2015b) (**Figure S1H**). As 20bp guides are markedly less tolerant of mismatches than longer ones for SaCas9 (type II-A SaCas9 and St1Cas9 share 37% identity), we favor the use of 20bp guides (Tycko et al., 2018). Hence, the flexibility of PAM recognition by St1Cas9 LMD-9 enhances its targeting capabilities. While recognition of an A-rich PAM may facilitate targeting A/T-rich regions of genomes, the targeting range of St1Cas9 in mammalian cells is less constrained than originally thought.

### Engineering St1Cas9 variants to expand its targeting range

Although not formally defined as the PAM at the time, the presence of a degenerate consensus sequence situated downstream of protospacers has been observed in strains of *S. thermophilus* almost 15 years ago (Bolotin et al., 2005). For example, inferred consensus PAM sequences for St1Cas9 from strains CNRZ1066 and LMG13811 are NNACAA(W) and NNGYAA(A) (where Y is C or T), respectively (Bolotin et al., 2005). Accordingly, the CRISPR1-Cas system of *S. thermophilus* strain LMG13811 transplanted in *E. coli* or reconstituted from purified components has been shown to target DNA using a NNGCAAA PAM (Chen et al., 2014).

At the protein level, the sequence of those St1Cas9 strain variants diverges mostly within the C-terminal wedge (WED) and PAM-interacting (PI) domains, implying that they have evolved to recognize distinct PAM sequences (**Figure S3A**). Since the PAM duplex is sandwiched between them, we tested whether swapping the WED and PI domains of St1Cas9 LMD-9 with the ones from LMG13811 and CNRZ1066 could reprogram PAM specificity (**Figure 2A**). While St1Cas9 LMD-9 could only target NNAGAA and NNGGAA PAMs, the hybrid constructs targeted with high efficacy and minimal cross reactivity NNGCAA and NNACAA PAMs, respectively (**Figure 2A**). Note that the same sgRNA architecture was used with all St1Cas9 variants in this setting which facilitates reprogramming.

**Figure 2.**
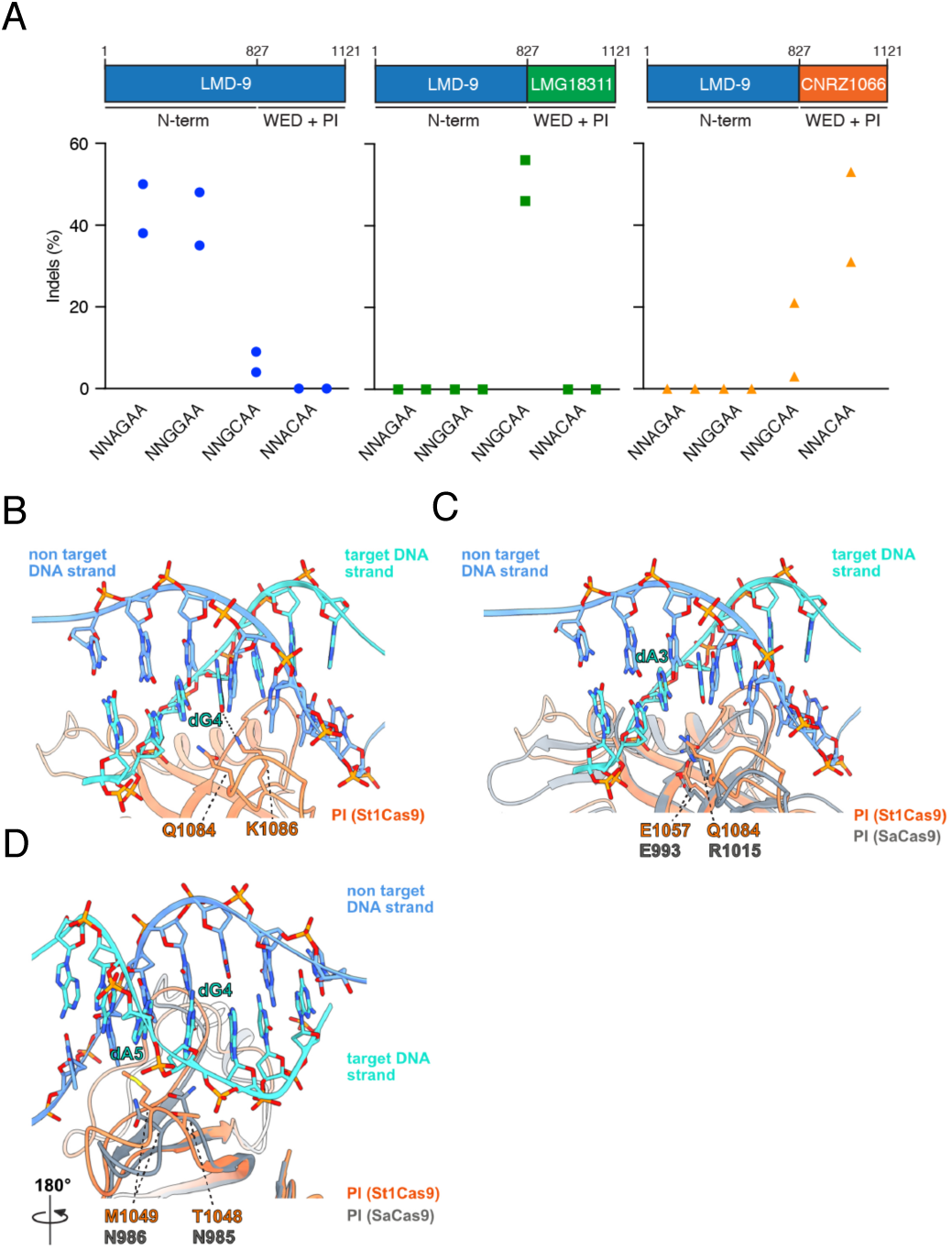
Structural basis for PAM specificity of engineered St1Cas9 variants with expanded targeting range. (A) Schematic representation of St1Cas9 hybrid proteins containing the N-terminal of LMD-9 and the C-terminal domains (WED + PI) of LMG18311 or CNRZ1066. To determine the activity of St1Cas9 variants programmed with sgRNAs targeting different PAMs, K562 cells were transiently transfected with single vector constructs driving expression of St1Cas9 and its sgRNA. Surveyor assays were performed 3 days later to determine the frequency of indels. An expression vector encoding EGFP (-) was used as a negative control. The experiment was performed twice and yielded equivalent results. (B) Close-up view of the 5’-GCAGAAA-3’ PAM bound to the St1Cas9 (DGCC7010) PI domain (PDB: 6RJD). The target (turquoise) and non-target (blue) strands are shown as sticks (the phosphate-sugar backbones are also shown as ribbons). The ribbon representation of the PI domain is orange. The hydrogen bonds between the side chain of St1Cas9 K1086 and the nucleobase of dG4 is shown as a dashed line. (C-D) The PI domains of St1Cas9 and SaCas9 (PDB: 5CZZ, grey ribbon) are superimposed. In (C), the St1Cas9 Q1084 and SaCas9 R1015 occupy the same position relative to the PAM (dA3). The St1Cas9 E1057 and SaCas9 E993 occupy the same position relative to St1Cas9 Q1084 and SaCas9 R1015, respectively. In (D), St1Cas9 T1048 and M1049 (substituted for N1048 and D1049 in some variants), superimpose onto SaCas9 N985 and N986 that specifies purines in positions 4 and 5 of the PAM. (See also **Figure S3**).

Sequence database mining using the “Search for PAMs by ALignment Of Targets” (SPAMALOT) tool (Chatterjee et al., 2018) predicts that even more diversity exists within CRISPR1-StCas9 systems and two additional groups represented by strains TH1477 and MTH17CL396 potentially target NNGAAA and NNAAAA PAMs, respectively (**Figure S3B**). Using the strategy described above, we were able to construct a highly active nuclease based on St1Cas9 TH1477 targeting the NNGAAA PAM (**Figure S3C**). These data highlight the modularity inherent to Cas9 enzymes and a simple strategy to further expand the targeting range of St1Cas9s. Currently, this set of four nucleases based on the St1Cas9 backbone can target five unique A-rich PAMs (NNAGAA, NNGGAA, NNGCAA, NNACAA, and NNGAAA). Tapping into the natural diversity found within *S. thermophilus* strains results in true reprograming towards a distinct PAM as opposed to relaxing specificity.

### Structural basis for St1Cas9s PAM specificity

We recently determined the structure of St1Cas9 (DGCC7710) bound to its sgRNA and to a target DNA containing a PAM to an overall resolution of 3.3 Å using single particle cryo-electron microscopy (*Cas9 allosteric inhibition by the anti-CRISPR protein AcrIIA6*, https://dx.doi.org/10.2139/ssrn.3403337). In this structure, the 5’-GCAGAAA-3’-containing PAM duplex is formed by seven Watson-Crick base pairs, and its major and minor grooves are sandwiched between the WED and PI domains. While the resolution of this structure prevents us from mapping all amino acid contacts with the PAM, the side chain of K1086 in the PI domain hydrogen bonds with the guanine at position 4 (NNA<u>G</u>_4_AA) (**Figure 2B**). Accordingly, St1Cas9 variants predicted by SPAMALOT (Chatterjee et al., 2018) to specify a guanine at this position contain K1086 (**Figures S3B** and **S3D**). There is only one type of substitution at that position, where K1086 is replaced by I1086 (**Figure S3D**). This set of variants, which includes LMG18311 (NNG<u>C</u>_<u>4</u>_AA), CNRZ1066 (NNA<u>C</u>_<u>4</u>_AA), and TH1477 (NNG<u>A</u>_<u>4</u>_AA) have lost the specificity for a guanine at position 4. At position 1084, the same type of analysis reveals that substitution of Q1084 for R1084 leads to the recognition of a guanine at position 3, as directly observed for LMG18311 (NN<u>G</u>_3_CAA) and TH1477 (NN<u>G</u>_3_AAA) (**Figures 2A** and **S3C**). Note that K1086 and R1084 are mutually exclusive and their co-occurrence would result in a steric clash (**Figure S3D**). In the structure of SaCas9 bound to 5’-TTG_3_AAT-3’ PAM, the third guanine is recognized by R1015 (Nishimasu et al., 2015). Structural comparison reveals that SaCas9 R1015 and St1Cas9 Q1084 occupy the same position relative to their PAMs (**Figure 2C**). In addition, SaCas9 R1015 is anchored via salt bridges to E993, a position equivalent to E1057 in St1Cas9 (**Figure 2C**). Thus, St1Cas9 variants with R1084 likely recognize guanine at position 3 in an analogous manner as SaCas9 does. Finally, a distinct set of amino acids surrounding positions 1048-1052 likely specify an adenine at position 4 in some variants as it is the case for TH1477 (NNG<u>A</u>_4_AA), and potentially, MTH17CL396 (NNA<u>A</u>_4_AA) (**Figure S3D**). T1048 and M1049 are replaced by N1048 and D1049 in those St1Cas9 variants and are predicted to occupy the same positions as N985 and N986 in SaCas9, the residues that specify purines at positions 4 and 5 in the NNGR_4_R_5_T PAM (where R is A or G) (Nishimasu et al., 2015) (**Figures 2D** and **S3D**). Structural comparison predicts that N1048 could directly contact the adenine in position 4, and that N1048 and D1049 would contact the adenine in positions 4 and 5, potentially via water-mediated hydrogen bonds as observed in SaCas9 (**Figure 2D**) (Nishimasu et al., 2015). Taken together, these observations provide a first glimpse at PAM recognition by St1Cas9 variants.

### Broadening the targeting scope of base editors using St1Cas9 variants

DNA base editors comprise fusions between a catalytically impaired Cas nuclease and a base modification enzyme that operates on single-stranded DNA (ssDNA) (Rees and Liu, 2018). Cytosine base editors (CBEs) convert a C• G base pair into a T• A using the APOBEC1 cytidine deaminase. Fusion of APOBEC1 to *Streptococcus pyogenes* Cas9 (SpCas9) D10A mutant (nickase) and two copies of the uracil DNA glycosylase inhibitor (UGI), resulted in the creation of the SpBE4max enzyme (Koblan et al., 2018). A limitation of the current base editing technology is that the protospacer adjacent motif (PAM) must be appropriately positioned relative to the target base to ensure efficient editing (Rees and Liu, 2018). Thus, there is a need to develop base editors with additional PAM compatibilities to increase the number of targetable bases in a genome. As such, SaCas9 has also been converted into a base editor to create SaBE4 (Rees and Liu, 2018). In an analogous manner, we have created St1BE4max by exchanging SpCas9 D10A for St1Cas9 D9A into the SpBE4max construct (Koblan et al., 2018). This created a potent CBE with novel targeting specificity due to the unique PAM of St1Cas9 (**Figure 3A**). Our data indicate that St1BE4max has an activity window similar to SaBE4, which is wider than SpBE4max, and sometimes extend upstream of the guide (Rees and Liu, 2018) (**Figure 3A**).

**Figure 3.**
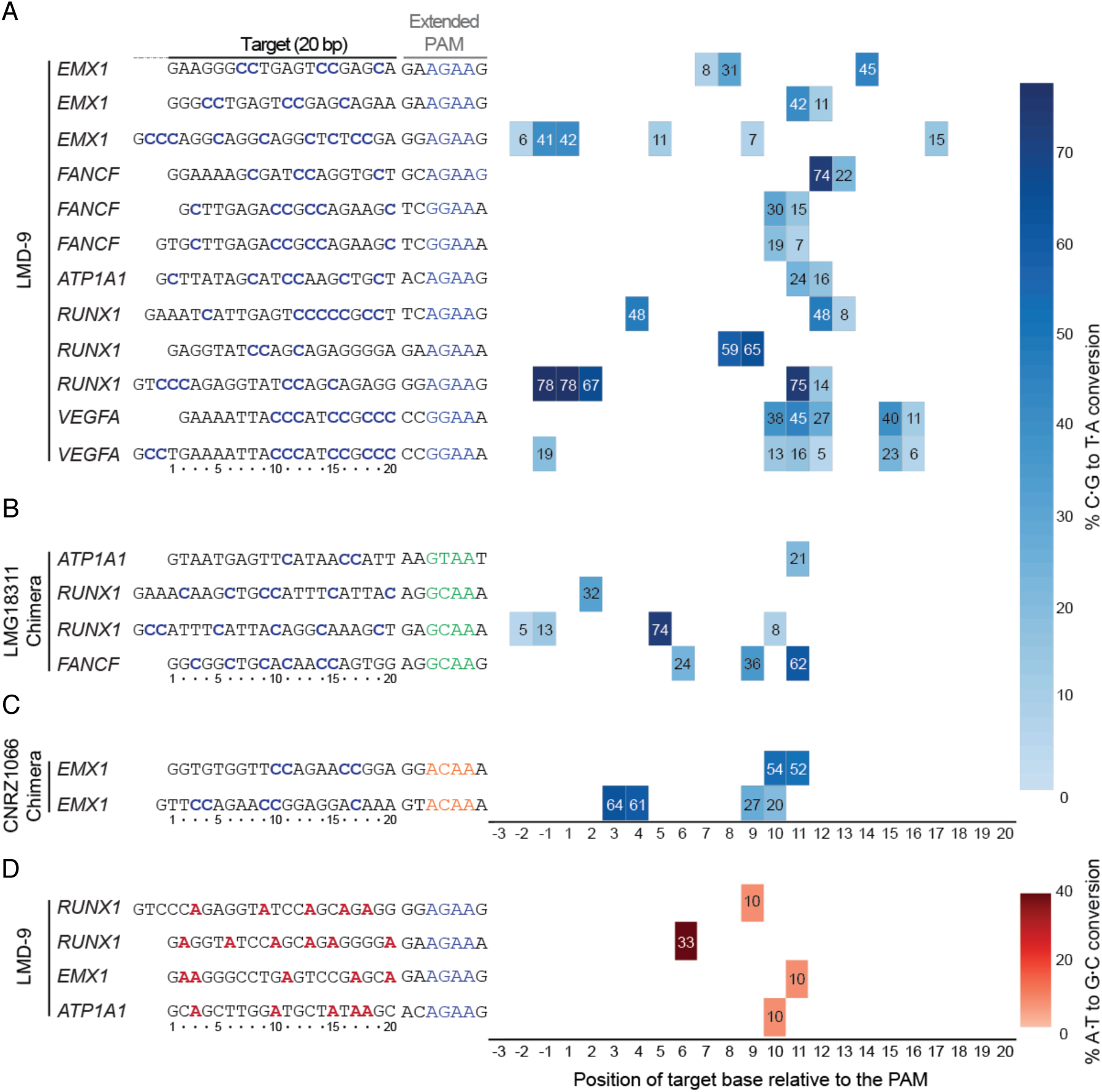
Broadening the targeting scope of base editors using St1Cas9 variants. (A) K562 cells were transiently transfected with single vector constructs driving expression of St1BE4max LMD-9 and its sgRNA. Genomic DNA was harvested 3 days later, and quantification of base editing was performed on PCR amplified target sites using EditR. The target sequence was defined as the 20 bases upstream of the PAM and numbered in decreasing order from the PAM. Sequence of the guides and related PAMs are shown with target Cs highlighted in blue. An expression vector encoding EGFP (-) was used as a negative control. (B-C) Same as (A) but using St1BE4max LMG18311 and CNRZ1066 chimeric proteins. (D) Same as (A) but using St1ABEmax LMD-9. Target adenines highlighted in red.

We then proceeded to test if St1Cas9 strain variants that display unique PAM preferences are also functional as CBEs. Indeed, LMG18311- and CNRZ1066-based St1BE4max are efficient base editors at NNGCAA and NNACAA PAMs, respectively (**Figures 3B-3C**). We also generated an adenine base editor (St1ABEmax) to mediate the conversion of A•T to G•C in genomic DNA. We observed moderate editing efficiencies of St1ABEmax, a phenomenon also observed for SaABEmax, indicating that the ABEmax architecture is not fully compatible with these shorter Cas9s (Huang et al., 2019) (**Figure 3D**). Nevertheless, these architectures can serve as a starting point for further improvements. Taken together, these data further demonstrate that St1Cas9 variants can be used as a scaffold to expand the targeting range of base editors.

### *In vivo* genome editing using St1Cas9

The small size of St1Cas9 makes it potentially permissive for packaging holo-St1Cas9 (St1Cas9 + sgRNA) into adeno-associated virus (AAV) vectors for *in vivo* delivery. To test the cleavage activity of St1Cas9 *in vivo*, we used the hereditary tyrosinemia type I (HT-I) mouse model, a disease caused by a deficiency of fumarylacetoacetate hydrolase (FAH), the last enzyme of the tyrosine catabolic pathway (<u>OMIM 276700</u>) (<u>Orphanet ORPHA:882</u>) (**Figure 4A**). *Fah-/-*mutant mice die as neonates with severe hepatic dysfunction and kidney damage due to the accumulation of toxic metabolites unless treated with nitisone (NTBC), a drug that inhibits 4-hydroxyphenylpyruvate dioxygenase (Hpd) upstream in the pathway (**Figure 4A**) (Grompe, 2017). Since genetic ablation of *Hpd* in mice can also prevent liver damage and lethality by creating a much milder HT-III phenotype (Endo et al., 1997; Pankowicz et al., 2016), we attempted to inactivate *Hpd* in our studies using St1Cas9.

**Figure 4.**
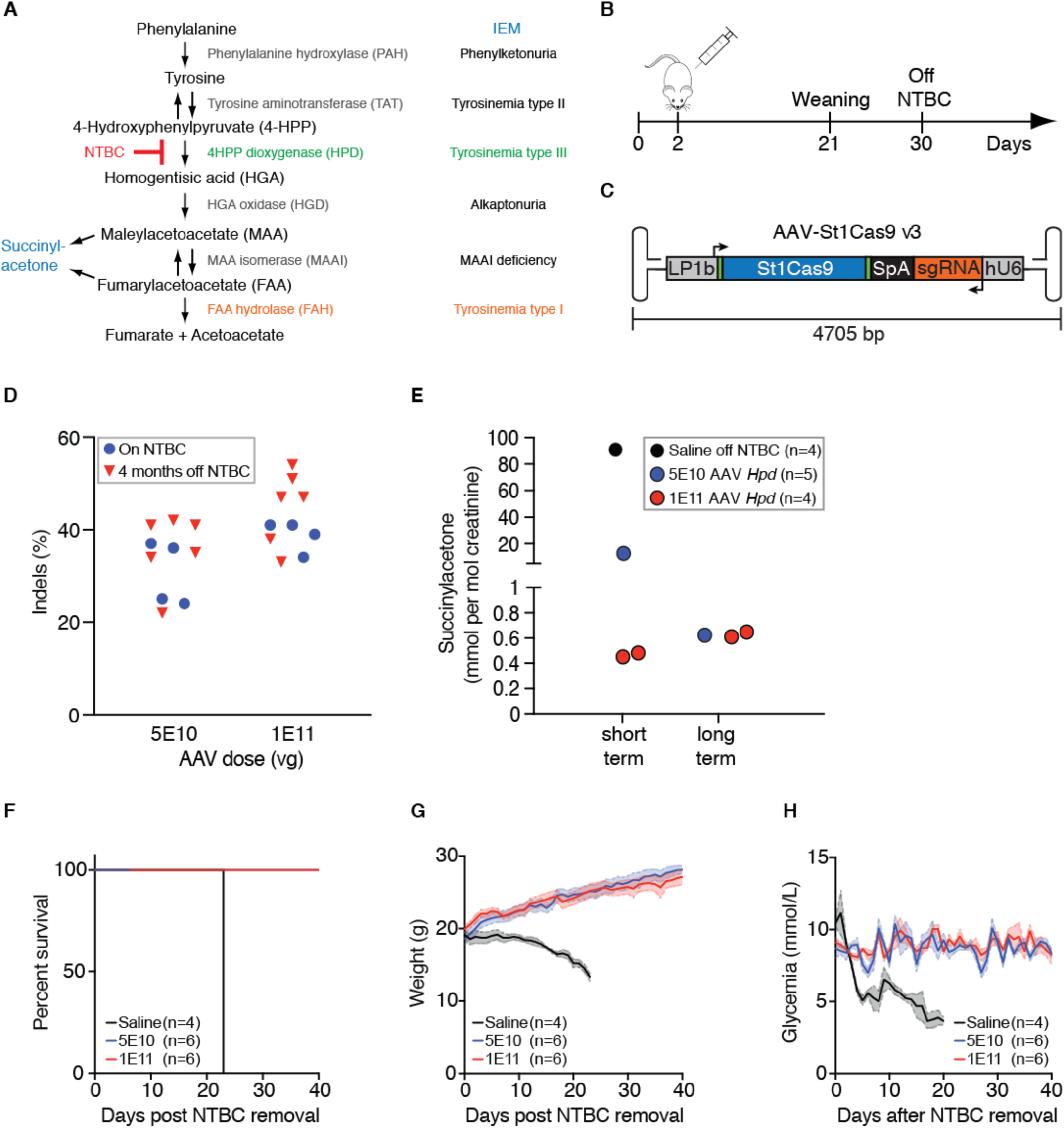
*In vivo* genome editing using St1Cas9. (A) The tyrosine degradation pathway and associated inborn errors of metabolism (IEM). (B) Experimental design. Neonatal (2 days old) *Fah*^*-/-*^ mice were injected with AAV8-St1Cas9 or saline into the retro-orbital sinus, weaned at 21 days, and NTBC was removed at 30 days of age. Mice off NTBC were killed when they lost 20% of their body weight. (C) Schematic representation of the AAV-St1Cas9 v3 vector. Annotated are the liver-specific promoter (LP1b) promoter, synthetic polyadenylation sequence (SpA) and hU6 promoter. Arrows indicate the direction of transcriptional unit. (D) Neonatal *Fah-/-*mice were injected with either 5E10 or 1E11 vector genomes (vg) of AAV8-St1Cas9 v3 targeting *Hpd* exon 13 and killed 28 days following injection or kept alive for phenotypic and metabolic studies for 4 months post NTBC removal. Genomic DNA was extracted from whole liver samples and the Surveyor assay was used to determine the frequency of indels. Each dot represents a different mouse. A mouse injected with saline (-) was used as a negative control. (E) Succinylacetone (SUAC) levels in urine from treated mice were determined 15 days (short term) or 4 months (long term) following NTBC removal. Samples were collected from the indicated treatment groups over a 24 hours period using metabolic cages. Number of mice per group/metabolic cage (n) and AAV doses (vg) is indicated. SUAC levels are undetectable in C57BL/6N (wild-type) mice. (F-H) Survival analysis, body weight, and glycemia following NTBC removal in treated mice. Body weight was measured daily and glycemia was monitored in non-fasted mice. Solid lines designate the mean and error bars are represented by shaded areas and denote s.e.m. (See also **Figure S4**).

To deliver holo-St1Cas9 to the liver, we generated a first set of AAV plasmids (AAV-St1Cas9 v1 and v2) containing a liver-specific promoter, sgRNA expression cassettes in opposite orientations and produced hepatotropic AAV serotype 8 (AAV8) vectors (Colella et al., 2018) (**Figures S4A-S4B**). We injected NTBC-treated *Fah*^*-/-*^ mice at day 2 of life into the retro-orbital sinus with these vectors and isolated total liver DNA at day 28 post injection in treated mice (**Figures 4B** and **S4A-S4B**). The titration showed that the degree of target editing at two different exons of *Hpd* was substantial and dependent on the dose of AAV8-St1Cas9 (**Figures S4A-S4B**). We then evaluated if alternative AAV-St1Cas9 expression cassettes could further improve cleavage efficacy *in vivo* while minimizing vector size. In our best performing design (AAV-St1Cas9 v3), we engineered a liver-specific promoter (LP1b) by combining elements from the human apolipoprotein E/C-I gene locus control region (*ApoE*-HCR), a modified human α1 antitrypsin promoter (h*AAT*), an SV40 intron, and used a synthetic polyadenylation signal element (**Figure 4C**) (McIntosh et al., 2013; Nathwani et al., 2006). These modifications increased cleavage efficacy markedly, especially at low AAV8 dose, and led to the creation of a vector of ∼4.7 kb in size which is optimal for viral particle packaging (**Figures 4C** and **4D**) (Colella et al., 2018). It is worth noting that as the genomic DNA was extracted from pieces of total livers, the effective activity is likely to be underestimated since hepatocytes make up 70% of the liver’s mass (Palaschak et al., 2019). Under the same experimental conditions, the levels of *in vivo* editing achieved with AAV-St1Cas9 v3 were comparable to the ones obtained using the gold standard AAV-SaCas9 system (**Figure S4C**) (Ran et al., 2015).

To test if AAV8-St1Cas9 v3 can achieve phenotypic correction *in vivo*, NTBC was withdrawn shortly after weaning in the remaining subset of treated mice. Systemic delivery via a single neonatal injection normalized the levels of excretion of succinylacetone (SUAC), a toxic metabolite and a diagnostic marker for HT-I (Grompe, 2017) (**Figure 4E**). Even at the lower vector dose (5E10), we observed delayed but near complete elimination of SUAC excretion 4 months following NTBC removal, which is likely due to the potent selective growth advantage of targeted hepatocytes that can extensively repopulate the diseased organ (Grompe, 2017) (**Figure 4E**). This is also reflected by the increased levels of indels detected in liver samples over the same period (**Figure 4D**). Consequently, treatment rescued lethality in all mice while saline-treated animals had to be killed after ∼3 weeks as they met the weight loss criterion (**Figure 4F**). Likewise, glycemia and weight loss were normalized in the treatment groups (**Figures 4G-4H**). Therefore, AAV8-mediated delivery of St1Cas9 in neonatal mice can result in efficient DNA cleavage, stable genetic modification, and phenotypic correction by rewiring a metabolic pathway through gene inactivation.

To corroborate these findings, we targeted the gene encoding phosphoenolpyruvate carboxykinase (Pck1 aka PEPCK), which plays a broad role in integrating hepatic energy metabolism and gluconeogenesis (Yang et al., 2009). Mice with a liver-specific deletion of the gene are viable but display an impaired response to fasting (She et al., 2000). Neonatal 2 days old C57BL/6N (wild-type) pups were injected with AAV8-St1Cas9 v3 targeting *Pck1* and at 6 weeks of age, they were fasted for 24 hours and killed for metabolic profiling and evaluating gene disruption efficacy (**Figures S4D-S4E**). Systemic delivery via a single neonatal injection resulted in substantial hepatic *Pck1* gene disruption (**Figure S4D)**. Plasma and hepatic triglyceride content were also markedly increased (**Figure S4E**). However, we found no change in circulating free fatty acid levels and hepatic glycogen stores were not depleted, suggesting that the observed phenotype may be intermediary to the one described in a prenatal hepatic knock-out model (**Figure S4E**) (She et al., 2000). We speculate that normal Pck1 in any non-targeted hepatocytes can partially compensate for the loss-of-function resulting from *in vivo* editing. Nevertheless, AAV8-mediated delivery of St1Cas9 in neonatal mice can efficiently disrupt the function of a key metabolic enzyme leading to clear and substantial phenotype *in vivo*. Collectively, these data support the notion that St1Cas9 can be engineered as a powerful tool for *in vivo* genome editing.

## DISCUSSION

Here we report that St1Cas9 can be harnessed for robust and efficient genome editing *in vitro* and *in vivo*, thereby expanding the CRISPR-Cas toolbox. We optimized this previously overlooked system to make nucleases, transcription activators, and base editors. We further validated its use in mice by demonstrating efficient rewiring, rescue, and creation of metabolic defects using all-in-one AAV vectors. Our work offers a comprehensive analysis and highlights novel fields of application for this CRISPR-Cas9 platform in mammalian cells (Kleinstiver et al., 2015b; Muller et al., 2016). St1Cas9 also functions efficiently for labeling of chromosomal loci in human cells and in mouse zygotes to create animal models (Fujii et al., 2016; Ma et al., 2015). In other systems, such as mycobacteria and the plant *Arabidopsis thaliana*, St1Cas9 is at least comparable to, and can even outperform, SpCas9 (Rock et al., 2017; Steinert et al., 2015). There is considerable interest in harnessing the diversity of Cas enzymes, but their implementation as genome editing tools is not a straightforward process (Chen et al., 2017; Ran et al., 2015; Zetsche et al., 2015). Nevertheless, orthologous Cas9 enzymes with different PAM requirements increase targeting flexibility and allow multiplexing for combinatorial genetic screens (Boettcher et al., 2018; Esvelt et al., 2013; Fonfara et al., 2014; Najm et al., 2018). This growing set of complementary and orthologous Cas systems could be adapted and combined for targeted knockout, transcriptional control, base editing, as well as “epigenome” editing (Komor et al., 2017).

Structure-guided and random mutagenesis have been combined to successfully reprogram the PI domain of Cas9s to alter its specificity towards a distinct sequence, but also to relax its specificity (for example NNGRRT would become NNNRRT or NGG would become NGN) (Hu et al., 2018; Kleinstiver et al., 2015a; Kleinstiver et al., 2015b; Nishimasu et al., 2018). As expected, relaxed PAM recognition typically decreases genome-wide specificity by increasing the number of off-targets (Kleinstiver et al., 2015a; Kleinstiver et al., 2019; Nishimasu et al., 2018). As a corollary, off-target sites are generally low in number for Cas9s with longer PAMs versus with shorter ones (Kleinstiver et al., 2015b; Kleinstiver et al., 2019; Tsai et al., 2015). This highlights an emerging connection between the length and complexity of the PAM versus the absolute specificity of a Cas protein that warrants further exploration (Muller et al., 2016). Interestingly, making use of the natural diversity within *S. thermophilus* strains allowed us to generate a suite of tools each with their unique PAM specificity to expand the targeting range of this CRISPR-based genome editing system. The engineering of CRISPR-Cas systems with unique PAM sequences is of utmost importance and should not be guided uniquely by the absolute targeting range (the total number of PAMs present in a genome) as the exact location of binding is most often key for genome editing. Applications such as, disruption of small genetic elements, allele-specific targeting, seamless gene correction via recombination, base editing, or gene correction via microhomology-mediated end joining require highly precise targeting (Canver et al., 2015; Gyorgy et al., 2019; Iyer et al., 2019; Rees and Liu, 2018). To achieve single nucleotide precision in targeting, a plethora of Cas9 orthologs harboring both wild-type and altered PAM specificities will be needed.

Recombinant AAV vectors are prime *in vivo* gene delivery vectors for non-proliferative tissues. However, a limitation in the therapeutic use of AAV is the loss of episomal vector genomes from actively dividing cells resulting in transient expression of therapeutic transgenes. Hence, the combination of genome editing technology with AAV-mediated delivery could lead to permanent genome modification and positive therapeutic outcome in young patients when tissues, such as the liver and retina, are still growing (Li et al., 2011; Yang et al., 2016). As a side benefit, the elimination of vector genomes would lead to transient nuclease expression in proliferating tissues that likely prevents accumulation of mutations at off-target sites (Li et al., 2011; Yang et al., 2016). In this perspective, the development of alternative *in vivo* genome editing platforms based on orthologous CRISPR-Cas systems would further increase the options available for therapeutic interventions.

## Supporting information

Protein_and_DNA_Sequences

## ACKNOWLEDGMENTS

This study was supported by grants from the Canadian Institutes of Health Research (CIHR) to M.L. and Y.D., and by the Banting Research Foundation to Y.D. S.M. acknowledges funding from NSERC. S.M. holds a Tier 1 Canada Research Chair in Bacteriophages. A.G. acknowledges funding from the French National Research Agency (ANR-18-CE11-0016-01). UCSF ChimeraX that was used for molecular graphics and analyses is developed by the Resource for Biocomputing, Visualization, and Informatics at the University of California, San Francisco, and receive support from NIH R01-GM129325 and P41-GM103311. Salary support was provided by the Fonds de la recherche du Québec-Santé (FRQS) to M.L. and Y.D. D.A. holds a Vanier Canada graduate scholarship. S.L. holds a Frederick Banting and Charles Best Canada graduate scholarship. A.D. and J-F.R. hold graduate training awards from the Fonds de la recherche du Québec-Santé (FRQS). We thank Marie-Ève Paquet and the skilled vector core facility staff at the Canadian neurophotonics platform for AAV8 production. Robert Tanguay provided the mouse model of HT-I, nitisone, expertise and support.

## AUTHOR CONTRIBUTIONS

Conceptualization, D.A., S.C., A.D., M.V., S.M., A.G., and Y.D.; Methodology, D.A., S.C., M.V., A.D., S.L., J.L., M.M., D.C., P.J.W., M.L., A.G., and Y.D.; Investigation, D.A., S.C., M.V., A.D., S.L., J-F. R., A.D., J.L., M.M., D.C.; Writing – Original Draft, Y.D.; Writing – Review and Editing, D.A., S.C., A.D., P.J.W., M.L., S.M., A.G., and Y.D.; Supervision, P.J.W., M.L., Y.D.; Funding Acquisition, M.L., S.M., A.G., and Y.D. All authors read and approved the final manuscript.

## METHODS

### Cell culture and transfection

K562 were obtained from the ATCC (CCL-243) and maintained at 37 °C under 5% CO_2_ in RPMI medium supplemented with 10% FBS, penicillin-streptomycin and GlutaMAX. Neuro-2a were obtained from the ATCC and maintained at 37 °C under 5% CO2 in DMEM medium supplemented with 10% FBS, penicillin-streptomycin and GlutaMAX. All cell lines are tested for absence of mycoplasma contamination. Cells (2E5 per transfection) were transfected using the Amaxa 4D-Nucleofector (Lonza) per manufacturer’s recommendations. Unless otherwise specified, 0.5 µg and 1 µg of single vector constructs driving the expression of both the sgRNA and St1Cas9 variants (nucleases or base editors) were used for transient transfections in Neuro-2a and K562 cells, respectively. K562 cell lines expressing St1Cas9 from the *AAVS1* safe harbor locus were generated as described (Agudelo et al., 2017; Dalvai et al., 2015). Briefly, simultaneous selection and cloning was performed for 10 days in methylcellulose-based semi-solid RPMI medium supplemented with 0.5 µg/ml puromycin starting 3 days post-transfection. Clones were picked and expanded in 96 wells for 3 days and transferred to 12-well plates for another 3 days before cells were harvested for western blot analysis.

### St1Cas9 strain variants

Sequences of St1Cas9 variants were retrieved from NCBI’s Identical Protein Groups (IPG) resource which contained 29 unique protein sequences at the time of analysis. Predicted PAM sequences for St1Cas9 LMD-9, LMG18311, CNRZ1066 were previously published (Bolotin et al., 2005; Horvath et al., 2008; Huang et al., 2019). Predictions for St1Cas9s related to TH1477 and MTH17CL396 originated from https://github.com/mitmedialab/SPAMALOT (Chatterjee et al., 2018). Alignments were performed using Clustal Omega (Sievers et al., 2011) in combination with the sequence alignment renderer ESPript 3 (Robert and Gouet, 2014).

### Structural analysis of PAM specificity

Coot and PISA were used to analyze the 3D structures (Krissinel and Henrick, 2007). UCSF ChimeraX was used to prepare the figures (Goddard et al., 2018).

### Genome editing vectors

Vectors for *in vitro* and *in vivo* genome editing with the CRISPR1-Cas9 (St1Cas9) system of *S. thermophilus* LMD-9 generated in this study are available from Addgene. Protein and DNA sequences for all St1Cas9 ORFs are available as supplemental items (Excel file). The mammalian expression vector for St1Cas9 (LMD-9) fused to SV40 NLS sequences at the N- and C-terminus (MSP1594_2x_NLS; Addgene plasmid #110625) was constructed from MSP1594 (Kleinstiver et al., 2015b) (Addgene plasmid #65775, a gift from Keith Joung). The U6-driven sgRNA expression cassettes for St1Cas9 (LMD-9) (v1, v2, v3) (St1Cas9_LMD-9_sgRNA_pUC19; Addgene plasmid #110627) were synthesized as gBlock gene fragments (Integrated DNA Technologies) and cloned into pUC19. BPK2301 (Kleinstiver et al., 2015b) (v0) (Addgene plasmid #65778, a gift from Keith Joung) was used to compare St1Cas9 sgRNA architectures. The single vector mammalian expression system containing a CAG promoter-driven St1Cas9 LMD-9 and its U6-driven sgRNA (U6_sgRNA_CAG_hSt1Cas9_LMD9; Addgene plasmid #110626) was built from the above-described plasmids. LMG18311, CNRZ1066, TH1477 C-terminal sequences were synthesized as gBlock gene fragments (Integrated DNA Technologies) and subcloned into U6_sgRNA_CAG_hSt1Cas9_LMD9 to produce the chimeric vectors.

Base editors were constructed into U6_sgRNA_CAG_hSt1Cas9_LMD9 (or the chimeric variants) using fragments derived from pCMV_BE4max_3xHA and pCMV_ABEmax_3xHA (Koblan et al., 2018) (Addgene plasmids #112096 and #112098, a gift from David Liu). Protein and DNA sequences for all St1Cas9 base editors are available as supplemental items (Excel file).

The single vector rAAV-St1Cas9 LMD-9 systems containing liver-specific promoters were assembled from the above-described components into a derivative of pX602 (Ran et al., 2015) (Addgene plasmid #61593, a gift from Feng Zhang) containing a deletion within the backbone to eliminate BsmBI restriction sites. The LP1b promoter was engineered by combining elements from previously described AAV expression cassettes (McIntosh et al., 2013; Nathwani et al., 2006). We deposited the most active version of this vector (v3) (pAAV_LP1B_St1Cas9_LMD-9_SpA_U6_sgRNA; Addgene plasmid #110624).

To establish clonal K562 cell lines constitutively expressing C-terminally tagged St1Cas9 under the control of an h*PGK1* promoter, the Cas9 ORF from MSP1594_2x_NLS was subcloned into AAVS1_Puro_PGK1_3xFLAG_Twin_Strep (Dalvai et al., 2015) (Addgene plasmid #68375).

The <u>CRISPOR</u> (Haeussler et al., 2016) web tool was used to design guide sequences against mouse and human targets St1Cas9 LMD-9. For St1Cas9 variants the guides were identified by manual inspection of target sequences. Guide sequences are available as supplemental items (Excel file).

### Surveyor nuclease, TIDE, and base editing assays

Genomic DNA from 2.5E5 cells was extracted with 250 µl of QuickExtract DNA extraction solution (Lucigen) per manufacturer’s recommendations. The various loci were amplified by 30 cycles of PCR using the primers described as supplemental items (Excel file). Assays were performed with the Surveyor mutation detection kit (Transgenomics) as described (Agudelo et al., 2017; Guschin et al., 2010). Samples were separated on 10% PAGE gels in TBE buffer. Gels were imaged using a ChemiDoc MP (Bio-Rad) system and quantifications were performed using the Image lab software (Bio-Rad). TIDE analysis was performed using a significance cut-off value for decomposition of p<0.001 (Brinkman et al., 2014). EditR (Kluesner et al., 2018) was used to quantify base editing from Sanger sequencing reads and all chromatograms are available as supplemental items (Excel file).

### Transcription reporter system

K562 cells were transfected with M_ST1n_VPR (0.25 µg) (aka dSt1Cas9-VPR) (Chavez et al., 2015) (Addgene plasmid #63799, a gift from George Church), M-tdTom-ST1 (0.25 µg) (Addgene #48678, a gift from George Church), and the indicated amounts of M-ST1-sgRNA (v0) (Addgene #48672, a gift from George Church) (Esvelt et al., 2013). Where indicated the sgRNA (v0) vector was exchanged for sgRNA (v1) (St1Cas9_LMD-9_sgRNA_pUC19; Addgene plasmid #110627) containing the same guide. Empty pUC19 vectors were used to normalize DNA concentration in all transfections. Fluorescence microscopy images were taken with an EVOS FL Cell Imaging System 3 days post-transfection. The intensity and the frequency of cells expressing tdTomato were assessed with a BD LSR II flow cytometer 3 days post-transfection.

### Recombinant adeno-associated virus production

Production of recombinant adeno-associated viral vectors was performed by the triple plasmid transfection method essentially as described (Gray et al., 2011). Briefly, HEK293T17 cells were transfected using polyethylenimine (PEI, Polysciences) with helper plasmid pxx-680 (A gift from R.J. Samulski), the rep/cap hybrid plasmid pAAV2/8 (A gift from James Wilson) and the rAAV vector plasmid. Twenty-four hours post-transfection, media was replaced with growth media without FBS, and cells were harvested 24 hours later. rAAV particles were extracted from cell extracts by freeze/thaw cycles and purified on a discontinuous iodixanol gradient. Virus were resuspended in PBS 320 mM NaCl + 5% D-sorbitol + 0.001% pluronic acid (F-68), aliquoted and stored at ^−^ 80°C. rAAV were titrated by qPCR (Roche) using SYBR green and ITR primers as described (Aurnhammer et al., 2012). The yields for all vectors varied between 1E13 and 2E13 vg/ml. The purity of the viral preparations was determined by SDS-PAGE analysis on a 10% stain free gel (Biorad) in Tris-Glycine-SDS buffer. ITR integrity was assessed following a BssHII digestion of the AAV plasmid. The vector core facility at the Canadian neurophotonics platform (molecular tools) produced the rAAV8s.

### Animal experiments (*Fah*^*-/-*^ mouse model)

*Fah*^*-/-*^ mice (Grompe et al., 1993) on a C57BL/6 genetic background were group-housed and fed a standard chow diet (Harlan #2018SX) with free access to food and water. *Fah*^*-/-*^ mice drinking water was supplemented with 7.5 mg (2-(2-nitro-4-trifluoromethylbenzoyl)-1,3-cyclohexanedione) (NTBC)/L and pH was adjusted to 7.0. Mice were exposed to a 12:12-h dark-light cycle and kept under an ambient temperature of 23 ± 1 °C. Animals were cared for and handled according to the *Canadian Guide for the Care and Use of Laboratory Animals.* The Université Laval Animal Care and Use Committee approved the procedures.

Two days old neonatal mice were injected intravenously in the retro-orbital sinus (Yardeni et al., 2011) with different doses of rAAV8 or saline in a total volume of 20 µL. Mice were weaned at 21 days of age and NTBC was removed 7 days later. Body weight and glycemia were monitored daily following NTBC removal. Mice were not fasted for measurement of glycemia, data collection occurred between 9-10 am. Animals were killed by cardiac puncture under anesthesia at predetermined time points or when weight loss reached 20% of body weight. Livers were snap frozen for downstream applications.

### Urine collection and succinylacetone quantification

Urine from groups of 3-4 mice was collected overnight in metabolic cages (Tecniplast) 15 days and 4 months after NTBC removal. Urine was centrifuged at 2000 rpm for 5 minutes, aliquoted and frozen at -80°C. Succinylacetone was quantified in urine samples by a sensitive method using gas chromatography–mass spectrometry (GC-MS) as previously described (Cyr et al., 2006). The biochemical genetics laboratory at the centre hospitalier universitaire de Sherbrooke performed the analyses.

### Animal experiments (C57BL/6)

Two days old neonatal *C57BL/6N* mice were injected intravenously in the retro-orbital sinus with 1E11 vector genome of rAAV8 targeting *Pck1* or saline in a total volume of 20 µL. At 6 weeks of age, mice were fasted for 24 hours and sacrificed by cardiac puncture under anaesthesia using syringes conditioned with EDTA. Liver sections were snap frozen and fixed in 4% PFA for downstream applications. Plasma was stored at -80°C for further biochemical analyses.

### Plasma triglycerides and NEFAs quantification

Plasma metabolites were assayed in duplicate using the following commercial kits; triglycerides (Thermo Fisher Scientific, TR22421), NEFAs (Wako, 999-34691, 995-34791, 991-34891, 993-35191, 276-76491).

### Hepatic triglyceride content extraction and measurement

Hepatic triglycerides extraction was previously described in detail (Caron et al., 2017). Briefly, liver sections (25-50 mg) were homogenized in a 2:1 chloroform:methanol mixture, combined with methanol, and centrifuged for 15 min at 3,000 rpm. 825 µl of supernatant was transferred to a new glass tube, chloroform and 0.73% NaCl was added, and the resulting mixture was centrifuged at 5,000 rpm for 3 min. The upper phase was discarded and the lower phase was washed 3 times with a 3/48/47 mixture of chloroform:methanol:NaCl (0.58%). The lower phase was then evaporated and resuspended in 1 ml of fresh isopropanol. Triglyceride levels were determined with a standard assay kit (Thermo Fisher Scientific, TR22421) according to the manufacturer’s instructions.

### Hepatic glycogen content extraction and measurement

Glycogen was measured in liver samples as described (Lo et al., 1970). Briefly, glycogen was extracted in 30% KOH saturated with Na_2_SO_4_, precipitated in 95% ethanol, and re-suspended in distilled H_2_O. Absorbance at 490 nm was measured in triplicates, after addition of phenol and H_2_SO_4_.

---

**Fig. S1.**
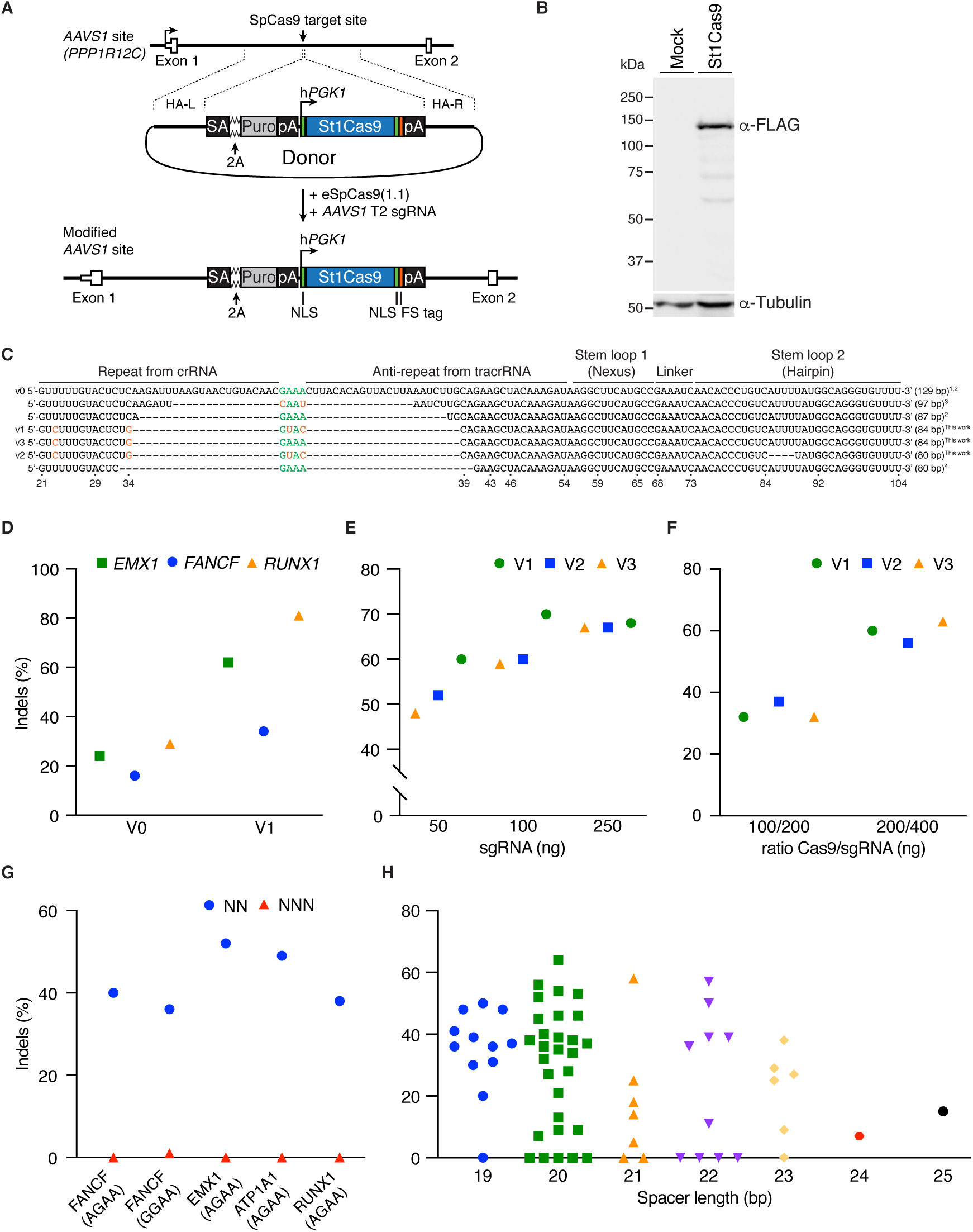
Related to Figure 1. Engineered CRISPR1-StCas9 system drives robust gene editing in mammalian cells. (A) Schematic representation of the targeted integration of tagged St1Cas9 to the *AAVS1* safe harbor locus. The donor construct and the locus following cDNA addition are displayed. The first two exons of the *PPP1R12C* gene are shown as open boxes. Also annotated are the locations of the splice acceptor site (SA), 2A self-cleaving peptide sequence (2A), puromycin resistance gene (Puro), polyadenylation sequence (pA), human phosphoglycerate kinase 1 promoter (h*PGK1*), nuclear localization signals (NLS), and 3xFLAG-2xSTREP tandem affinity tag (Tag), homology arms left and right (HA-L, HA-R) are respectively 800 and 840 bp. (B) Western blots showing St1Cas9-tag protein expression in a K562 clone and in cells expressing only the tag (Mock). The FLAG M2 antibody was used to detect Cas9 and the tubulin antibody was used as a loading control. (C) Alignment of previously described sgRNA sequences for St1Cas9 as well as novel designs from the present study (v1, v2, v3). ^1^Kleinstiver, Nature 2015b, ^2^Esvelt, Nat Methods 2013, ^3^Muller, Mol Ther 2016, ^4^Ran, Nature 2015. (D) K562 cells were transiently transfected with a St1Cas9 LMD-9 expression vector (0.5μg) in addition to the indicated sgRNA expression plasmids (0.8μg) and Surveyor assays were performed 3 days later to determine the frequency of indels at the 3 specified targets. (E) K562 cells stably expressing St1Cas9 LMD-9 were transfected with indicated sgRNA expression vectors targeting *EMX1* at increasing doses and TIDE assays were performed 3 days later to determine the frequency of indels. (F) K562 cells were transiently transfected with a St1Cas9 LMD-9 expression vector (100 or 200 ng) in addition to the indicated sgRNA expression plasmids (200 and 400 ng) targeting *EMX1*. Indel frequency was determined as in (D). (G) sgRNAs specifying cleavage by St1Cas9 LMD-9 at indicated PAMs with an NN linker were modified to test their functionality with an NNN linker at the specified targets. Cleavage activity was determined as in (D). (H) Cleavage activity of sgRNAs shown in Figure 1C ordered by guide length.

**Fig. S2.**
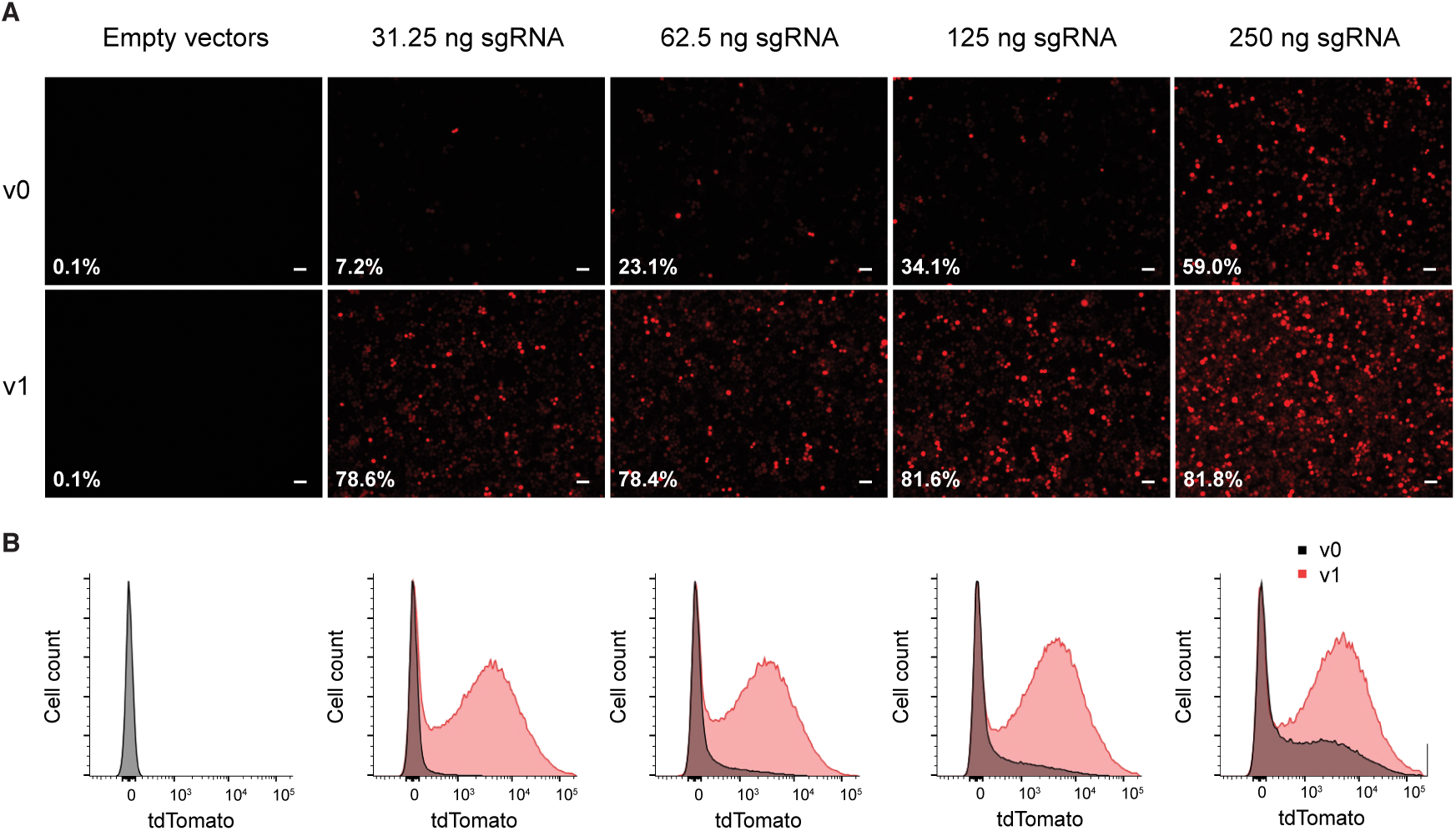
Related to Figure 1. Engineered CRISPR1-StCas9 sgRNA improves dSt1Cas9-VPR transcriptional activation in human cells. (A) K562 cells were transfected with the tdTomato transcriptional reporter along with dSt1Cas9-VPR and the targeting sgRNAs (v0 or v1) at increasing doses. Fluorescence microscopy images were taken 3 days post-transfection. Scale bars represent 50 µm. (B) Transcriptional activation was quantified by FACS 3 days post-transfection. % of tdTomato-positive cells are indicated in panel (A).

**Fig. S3.**
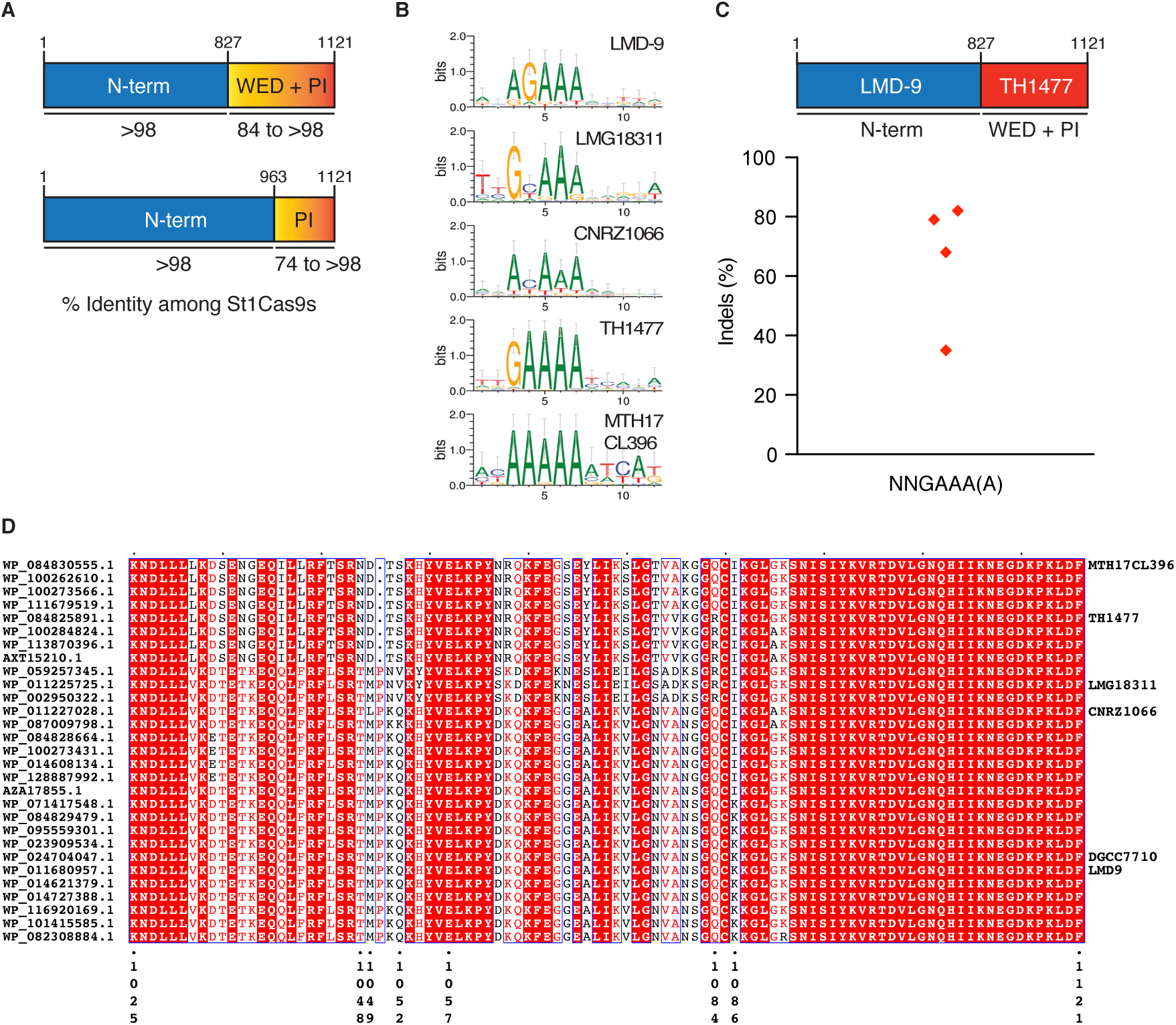
Related to Figure 2. St1Cas9 strain variants display unique PAM specificities. (A) Percentage of identity within the N-terminal versus the WED and PI domains of St1Cas9s. Amino acid sequence alignment of St1Cas9 respective domains from all 29 variants available in NCBI’s Identical Protein Groups (IPG) resource were performed with Clustal Omega. (B) Sequence logos of predicted PAMs for the indicated St1Cas9 “families” using SPAMALOT. (C) Schematic representation of St1Cas9 hybrid protein containing the N-terminal of LMD-9 and the C-terminal domains (WED + PI) of TH1477. K562 cells were transiently transfected with single vector constructs driving expression of St1Cas9 and its sgRNA. Surveyor assays were performed 3 days later to determine the frequency of indels. Each lozenge represents a different target. An expression vector encoding EGFP (-) was used as a negative control. (D) Amino acid sequence alignment of St1Cas9 CTD domains from all 29 variants available in NCBI’s Identical Protein Groups (IPG) resource. Alignment was performed with Clustal Omega and rendered using ESPript 3.

**Fig. S4.**
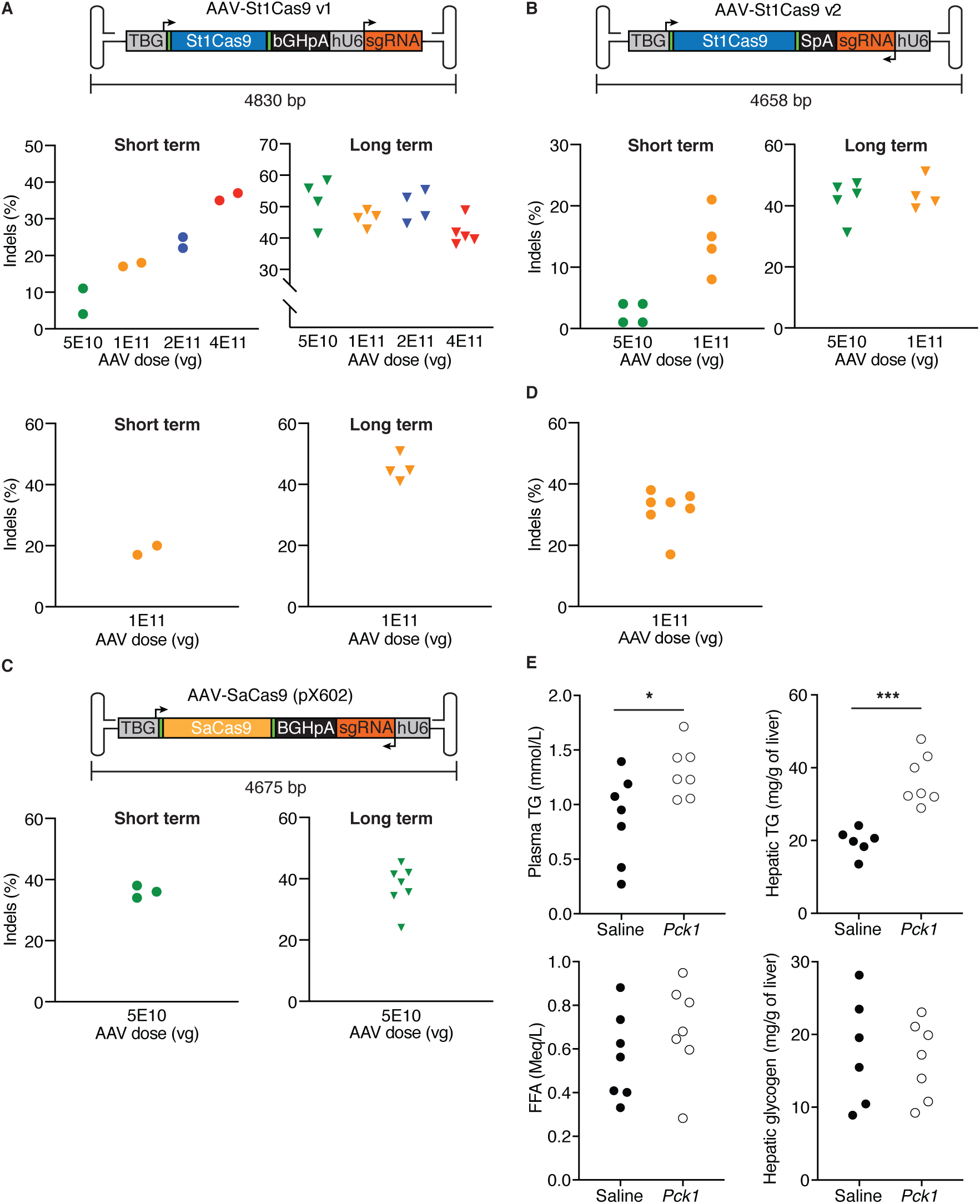
Related to Figure 4. *In vivo* genome editing using St1Cas9. (A) Schematic representation of the St1Cas9 AAV v1 vector. Human thyroxine binding globulin (TBG) promoter, bovine growth hormone polyadenylation sequence (BGHpA) and hU6 promoter are shown. Arrows indicate the direction of transcriptional unit. Neonatal *Fah*^*-/-*^mice were injected into the retro-orbital sinus with either 5E10, 1E11, 2E11 or 4E11 vector genomes (vg) of AAV8-St1Cas9 v1 targeting *Hpd* exon 13 (G5) (Top) or 1E11 vector genomes (vg) of AAV8-St1Cas9 v1 targeting *Hpd* exon 8 (G2) (Bot). Mice were killed 28 days following injection (short term) or kept alive for 8 months (long term) post NTBC removal. Genomic DNA was extracted from whole liver samples and the Surveyor assay (short term) or TIDE assay (long term) were used to determine the frequency of indels. Each dot represents a different mouse. A mouse injected with saline (-) was used as a negative control. (B) Same as in (A) but mice were injected with either 5E10 or 1E11 vector genomes (vg) of AAV8-St1Cas9 v2 targeting *Hpd* exon 13 (G5). (C) Same as in (A) but using 5E10 vector genomes (vg) of AAV8-SaCas9 targeting exon 7 of *Hpd*. Mice were killed after 12 months for the long-term condition. (D) Neonatal C57BL/6N mice were injected into the retro-orbital sinus with 1E11 vector genomes (vg) of AAV8-St1Cas9 v3 targeting exon 5 of *Pck1*, weaned at 21 days and fasted for 24 hours at 6 weeks of age before sacrifice. Genomic DNA was extracted from whole liver samples and the Surveyor assay was used to determine the frequency. Each dot represents a different mouse. A mouse injected with saline (-) was used as a negative control. (E) Plasma triglyceride (Top left) and free fatty acids (Bot left) levels were measured after 24 hours of fasting in mice injected with saline or AAV8-St1Cas9 v3. Hepatic triglycerides (Top right) and Hepatic glycogen (Bot right) were extracted and quantified. Each dot represents a different animal. * p < 0.05, *** p < 0.0005 according to Student’s test.

## REFERENCES

Agudelo, D., Duringer, A., Bozoyan, L., Huard, C.C., Carter, S., Loehr, J., Synodinou, D., Drouin, M., Salsman, J., Dellaire, G., et al. (2017). Marker-free coselection for CRISPR-driven genome editing in human cells. Nat Methods 14, 615–620.

Aurnhammer, C., Haase, M., Muether, N., Hausl, M., Rauschhuber, C., Huber, I., Nitschko, H., Busch, U., Sing, A., Ehrhardt, A., et al. (2012). Universal real-time PCR for the detection and quantification of adeno-associated virus serotype 2-derived inverted terminal repeat sequences. Hum Gene Ther Methods 23, 18–28.

Barrangou, R., and Horvath, P. (2017). A decade of discovery: CRISPR functions and applications. Nat Microbiol 2, 17092.

Boettcher, M., Tian, R., Blau, J.A., Markegard, E., Wagner, R.T., Wu, D., Mo, X., Biton, A., Zaitlen, N., Fu, H., et al. (2018). Dual gene activation and knockout screen reveals directional dependencies in genetic networks. Nat Biotechnol 36, 170–178.

Bolotin, A., Quinquis, B., Sorokin, A., and Ehrlich, S.D. (2005). Clustered regularly interspaced short palindrome repeats (CRISPRs) have spacers of extrachromosomal origin. Microbiology 151, 2551–2561.

Briner, A.E., Donohoue, P.D., Gomaa, A.A., Selle, K., Slorach, E.M., Nye, C.H., Haurwitz, R.E., Beisel, C.L., May, A.P., and Barrangou, R. (2014). Guide RNA functional modules direct Cas9 activity and orthogonality. Mol Cell 56, 333–339.

Brinkman, E.K., Chen, T., Amendola, M., and van Steensel, B. (2014). Easy quantitative assessment of genome editing by sequence trace decomposition. Nucleic Acids Res 42, e168.

Canver, M.C., Smith, E.C., Sher, F., Pinello, L., Sanjana, N.E., Shalem, O., Chen, D.D., Schupp, P.G., Vinjamur, D.S., Garcia, S.P., et al. (2015). BCL11A enhancer dissection by Cas9-mediated in situ saturating mutagenesis. Nature 527, 192–197.

Caron, A., Mouchiroud, M., Gautier, N., Labbe, S.M., Villot, R., Turcotte, L., Secco, B., Lamoureux, G., Shum, M., Gelinas, Y., et al. (2017). Loss of hepatic DEPTOR alters the metabolic transition to fasting. Mol Metab 6, 447–458.

Chari, R., Mali, P., Moosburner, M., and Church, G.M. (2015). Unraveling CRISPR-Cas9 genome engineering parameters via a library-on-library approach. Nat Methods 12, 823–826.

Chatterjee, P., Jakimo, N., and Jacobson, J.M. (2018). Minimal PAM specificity of a highly similar SpCas9 ortholog. Sci Adv 4, eaau0766.

Chavez, A., Scheiman, J., Vora, S., Pruitt, B.W., Tuttle, M., E, P.R.I., Lin, S., Kiani, S., Guzman, C.D., Wiegand, D.J., et al. (2015). Highly efficient Cas9-mediated transcriptional programming. Nat Methods 12, 326–328.

Chen, F., Ding, X., Feng, Y., Seebeck, T., Jiang, Y., and Davis, G.D. (2017). Targeted activation of diverse CRISPR-Cas systems for mammalian genome editing via proximal CRISPR targeting. Nat Commun 8, 14958.

Chen, H., Choi, J., and Bailey, S. (2014). Cut site selection by the two nuclease domains of the Cas9 RNA-guided endonuclease. J Biol Chem 289, 13284–13294.

Colella, P., Ronzitti, G., and Mingozzi, F. (2018). Emerging Issues in AAV-Mediated In Vivo Gene Therapy. Mol Ther Methods Clin Dev 8, 87–104.

Cong, L., Ran, F.A., Cox, D., Lin, S., Barretto, R., Habib, N., Hsu, P.D., Wu, X., Jiang, W., Marraffini, L.A., et al. (2013). Multiplex genome engineering using CRISPR/Cas systems. Science 339, 819–823.

Cyr, D., Giguere, R., Villain, G., Lemieux, B., and Drouin, R. (2006). A GC/MS validated method for the nanomolar range determination of succinylacetone in amniotic fluid and plasma: an analytical tool for tyrosinemia type I. J Chromatogr B Analyt Technol Biomed Life Sci 832, 24–29.

Dalvai, M., Loehr, J., Jacquet, K., Huard, C.C., Roques, C., Herst, P., Cote, J., and Doyon, Y. (2015). A Scalable Genome-Editing-Based Approach for Mapping Multiprotein Complexes in Human Cells. Cell Rep 13, 621–633.

Deveau, H., Barrangou, R., Garneau, J.E., Labonte, J., Fremaux, C., Boyaval, P., Romero, D.A., Horvath, P., and Moineau, S. (2008). Phage response to CRISPR-encoded resistance in Streptococcus thermophilus. J Bacteriol 190, 1390–1400.

Edraki, A., Mir, A., Ibraheim, R., Gainetdinov, I., Yoon, Y., Song, C.Q., Cao, Y., Gallant, J., Xue, W., Rivera-Perez, J.A., et al. (2019). A Compact, High-Accuracy Cas9 with a Dinucleotide PAM for In Vivo Genome Editing. Mol Cell 73, 714–726 e714.

Endo, F., Kubo, S., Awata, H., Kiwaki, K., Katoh, H., Kanegae, Y., Saito, I., Miyazaki, J., Yamamoto, T., Jakobs, C., et al. (1997). Complete rescue of lethal albino c14CoS mice by null mutation of 4-hydroxyphenylpyruvate dioxygenase and induction of apoptosis of hepatocytes in these mice by in vivo retrieval of the tyrosine catabolic pathway. J Biol Chem 272, 24426–24432.

Esvelt, K.M., Mali, P., Braff, J.L., Moosburner, M., Yaung, S.J., and Church, G.M. (2013). Orthogonal Cas9 proteins for RNA-guided gene regulation and editing. Nat Methods 10, 1116–1121.

Fonfara, I., Le Rhun, A., Chylinski, K., Makarova, K.S., Lecrivain, A.L., Bzdrenga, J., Koonin, E.V., and Charpentier, E. (2014). Phylogeny of Cas9 determines functional exchangeability of dual-RNA and Cas9 among orthologous type II CRISPR-Cas systems. Nucleic Acids Res 42, 2577–2590.

Fujii, W., Kakuta, S., Yoshioka, S., Kyuwa, S., Sugiura, K., and Naito, K. (2016). Zygote-mediated generation of genome-modified mice using Streptococcus thermophilus 1-derived CRISPR/Cas system. Biochem Biophys Res Commun 477, 473–476.

Goddard, T.D., Huang, C.C., Meng, E.C., Pettersen, E.F., Couch, G.S., Morris, J.H., and Ferrin, T.E. (2018). UCSF ChimeraX: Meeting modern challenges in visualization and analysis. Protein Sci 27, 14–25.

Gray, S.J., Choi, V.W., Asokan, A., Haberman, R.A., McCown, T.J., and Samulski, R.J. (2011). Production of recombinant adeno-associated viral vectors and use in in vitro and in vivo administration. Curr Protoc Neurosci Chapter 4, Unit 4 17.

Grompe, M. (2017). Fah Knockout Animals as Models for Therapeutic Liver Repopulation. Adv Exp Med Biol 959, 215–230.

Grompe, M., al-Dhalimy, M., Finegold, M., Ou, C.N., Burlingame, T., Kennaway, N.G., and Soriano, P. (1993). Loss of fumarylacetoacetate hydrolase is responsible for the neonatal hepatic dysfunction phenotype of lethal albino mice. Genes Dev 7, 2298–2307.

Guschin, D.Y., Waite, A.J., Katibah, G.E., Miller, J.C., Holmes, M.C., and Rebar, E.J. (2010). A rapid and general assay for monitoring endogenous gene modification. Methods Mol Biol 649, 247–256.

Gyorgy, B., Nist-Lund, C., Pan, B., Asai, Y., Karavitaki, K.D., Kleinstiver, B.P., Garcia, S.P., Zaborowski, M.P., Solanes, P., Spataro, S., et al. (2019). Allele-specific gene editing prevents deafness in a model of dominant progressive hearing loss. Nat Med.

Haeussler, M., Schonig, K., Eckert, H., Eschstruth, A., Mianne, J., Renaud, J.B., Schneider-Maunoury, S., Shkumatava, A., Teboul, L., Kent, J., et al. (2016). Evaluation of off-target and on-target scoring algorithms and integration into the guide RNA selection tool CRISPOR. Genome Biol 17, 148.

Hille, F., Richter, H., Wong, S.P., Bratovic, M., Ressel, S., and Charpentier, E. (2018). The Biology of CRISPR-Cas: Backward and Forward. Cell 172, 1239–1259.

Horvath, P., Romero, D.A., Coute-Monvoisin, A.C., Richards, M., Deveau, H., Moineau, S., Boyaval, P., Fremaux, C., and Barrangou, R. (2008). Diversity, activity, and evolution of CRISPR loci in Streptococcus thermophilus. J Bacteriol 190, 1401–1412.

Hu, J.H., Miller, S.M., Geurts, M.H., Tang, W., Chen, L., Sun, N., Zeina, C.M., Gao, X., Rees, H.A., Lin, Z., et al. (2018). Evolved Cas9 variants with broad PAM compatibility and high DNA specificity. Nature 556, 57–63.

Huang, T.P., Zhao, K.T., Miller, S.M., Gaudelli, N.M., Oakes, B.L., Fellmann, C., Savage, D.F., and Liu, D.R. (2019). Circularly permuted and PAM-modified Cas9 variants broaden the targeting scope of base editors. Nat Biotechnol 37, 626–631.

Hynes, A.P., Rousseau, G.M., Agudelo, D., Goulet, A., Amigues, B., Loehr, J., Romero, D.A., Fremaux, C., Horvath, P., Doyon, Y., et al. (2018). Widespread anti-CRISPR proteins in virulent bacteriophages inhibit a range of Cas9 proteins. Nat Commun 9, 2919.

Ibraheim, R., Song, C.Q., Mir, A., Amrani, N., Xue, W., and Sontheimer, E.J. (2018). All-in-one adeno-associated virus delivery and genome editing by Neisseria meningitidis Cas9 in vivo. Genome Biol 19, 137.

Iyer, S., Suresh, S., Guo, D., Daman, K., Chen, J.C.J., Liu, P., Zieger, M., Luk, K., Roscoe, B.P., Mueller, C., et al. (2019). Precise therapeutic gene correction by a simple nuclease-induced double-stranded break. Nature 568, 561–565.

Jinek, M., Chylinski, K., Fonfara, I., Hauer, M., Doudna, J.A., and Charpentier, E. (2012). A programmable dual-RNA-guided DNA endonuclease in adaptive bacterial immunity. Science 337, 816–821.

Kim, E., Koo, T., Park, S.W., Kim, D., Kim, K., Cho, H.Y., Song, D.W., Lee, K.J., Jung, M.H., Kim, S., et al. (2017). In vivo genome editing with a small Cas9 orthologue derived from Campylobacter jejuni. Nat Commun 8, 14500.

Kleinstiver, B.P., Prew, M.S., Tsai, S.Q., Nguyen, N.T., Topkar, V.V., Zheng, Z., and Joung, J.K. (2015a). Broadening the targeting range of Staphylococcus aureus CRISPR-Cas9 by modifying PAM recognition. Nat Biotechnol 33, 1293–1298.

Kleinstiver, B.P., Prew, M.S., Tsai, S.Q., Topkar, V.V., Nguyen, N.T., Zheng, Z., Gonzales, A.P., Li, Z., Peterson, R.T., Yeh, J.R., et al. (2015b). Engineered CRISPR-Cas9 nucleases with altered PAM specificities. Nature 523, 481–485.

Kleinstiver, B.P., Sousa, A.A., Walton, R.T., Tak, Y.E., Hsu, J.Y., Clement, K., Welch, M.M., Horng, J.E., Malagon-Lopez, J., Scarfo, I., et al. (2019). Engineered CRISPR-Cas12a variants with increased activities and improved targeting ranges for gene, epigenetic and base editing. Nat Biotechnol.

Kluesner, M.G., Nedveck, D.A., Lahr, W.S., Garbe, J.R., Abrahante, J.E., Webber, B.R., and Moriarity, B.S. (2018). EditR: A Method to Quantify Base Editing from Sanger Sequencing. CRISPR J 1, 239–250.

Koblan, L.W., Doman, J.L., Wilson, C., Levy, J.M., Tay, T., Newby, G.A., Maianti, J.P., Raguram, A., and Liu, D.R. (2018). Improving cytidine and adenine base editors by expression optimization and ancestral reconstruction. Nat Biotechnol 36, 843–846.

Komor, A.C., Badran, A.H., and Liu, D.R. (2017). CRISPR-Based Technologies for the Manipulation of Eukaryotic Genomes. Cell 169, 559.

Koonin, E.V., Makarova, K.S., and Zhang, F. (2017). Diversity, classification and evolution of CRISPR-Cas systems. Curr Opin Microbiol 37, 67–78.

Krissinel, E., and Henrick, K. (2007). Inference of macromolecular assemblies from crystalline state. J Mol Biol 372, 774–797.

Lau, C.H., and Suh, Y. (2017). In vivo genome editing in animals using AAV-CRISPR system: applications to translational research of human disease. F1000Res 6, 2153.

Leenay, R.T., Maksimchuk, K.R., Slotkowski, R.A., Agrawal, R.N., Gomaa, A.A., Briner, A.E., Barrangou, R., and Beisel, C.L. (2016). Identifying and Visualizing Functional PAM Diversity across CRISPR-Cas Systems. Mol Cell 62, 137–147.

Li, H., Haurigot, V., Doyon, Y., Li, T., Wong, S.Y., Bhagwat, A.S., Malani, N., Anguela, X.M., Sharma, R., Ivanciu, L., et al. (2011). In vivo genome editing restores haemostasis in a mouse model of haemophilia. Nature 475, 217–221.

Lo, S., Russell, J.C., and Taylor, A.W. (1970). Determination of glycogen in small tissue samples. J Appl Physiol 28, 234–236.

Ma, H., Naseri, A., Reyes-Gutierrez, P., Wolfe, S.A., Zhang, S., and Pederson, T. (2015). Multicolor CRISPR labeling of chromosomal loci in human cells. Proc Natl Acad Sci U S A 112, 3002–3007.

Makarova, K.S., Wolf, Y.I., and Koonin, E.V. (2018). Classification and Nomenclature of CRISPR-Cas Systems: Where from Here? CRISPR J 1, 325–336.

McIntosh, J., Lenting, P.J., Rosales, C., Lee, D., Rabbanian, S., Raj, D., Patel, N., Tuddenham, E.G., Christophe, O.D., McVey, J.H., et al. (2013). Therapeutic levels of FVIII following a single peripheral vein administration of rAAV vector encoding a novel human factor VIII variant. Blood 121, 3335–3344.

Mir, A., Edraki, A., Lee, J., and Sontheimer, E.J. (2018). Type II-C CRISPR-Cas9 Biology, Mechanism, and Application. ACS Chem Biol 13, 357–365.

Muller, M., Lee, C.M., Gasiunas, G., Davis, T.H., Cradick, T.J., Siksnys, V., Bao, G., Cathomen, T., and Mussolino, C. (2016). Streptococcus thermophilus CRISPR-Cas9 Systems Enable Specific Editing of the Human Genome. Mol Ther 24, 636–644.

Najm, F.J., Strand, C., Donovan, K.F., Hegde, M., Sanson, K.R., Vaimberg, E.W., Sullender, M.E., Hartenian, E., Kalani, Z., Fusi, N., et al. (2018). Orthologous CRISPR-Cas9 enzymes for combinatorial genetic screens. Nat Biotechnol 36, 179–189.

Nathwani, A.C., Gray, J.T., Ng, C.Y., Zhou, J., Spence, Y., Waddington, S.N., Tuddenham, E.G., Kemball-Cook, G., McIntosh, J., Boon-Spijker, M., et al. (2006). Self-complementary adeno-associated virus vectors containing a novel liver-specific human factor IX expression cassette enable highly efficient transduction of murine and nonhuman primate liver. Blood 107, 2653–2661.

Nishimasu, H., Cong, L., Yan, W.X., Ran, F.A., Zetsche, B., Li, Y., Kurabayashi, A., Ishitani, R., Zhang, F., and Nureki, O. (2015). Crystal Structure of Staphylococcus aureus Cas9. Cell 162, 1113–1126.

Nishimasu, H., Shi, X., Ishiguro, S., Gao, L., Hirano, S., Okazaki, S., Noda, T., Abudayyeh, O.O., Gootenberg, J.S., Mori, H., et al. (2018). Engineered CRISPR-Cas9 nuclease with expanded targeting space. Science 361, 1259–1262.

Palaschak, B., Herzog, R.W., and Markusic, D.M. (2019). AAV-Mediated Gene Delivery to the Liver: Overview of Current Technologies and Methods. Methods Mol Biol 1950, 333–360.

Pankowicz, F.P., Barzi, M., Legras, X., Hubert, L., Mi, T., Tomolonis, J.A., Ravishankar, M., Sun, Q., Yang, D., Borowiak, M., et al. (2016). Reprogramming metabolic pathways in vivo with CRISPR/Cas9 genome editing to treat hereditary tyrosinaemia. Nat Commun 7, 12642.

Ran, F.A., Cong, L., Yan, W.X., Scott, D.A., Gootenberg, J.S., Kriz, A.J., Zetsche, B., Shalem, O., Wu, X., Makarova, K.S., et al. (2015). In vivo genome editing using Staphylococcus aureus Cas9. Nature 520, 186–191.

Rees, H.A., and Liu, D.R. (2018). Base editing: precision chemistry on the genome and transcriptome of living cells. Nat Rev Genet 19, 770–788.

Robert, X., and Gouet, P. (2014). Deciphering key features in protein structures with the new ENDscript server. Nucleic Acids Res 42, W320–324.

Rock, J.M., Hopkins, F.F., Chavez, A., Diallo, M., Chase, M.R., Gerrick, E.R., Pritchard, J.R., Church, G.M., Rubin, E.J., Sassetti, C.M., et al. (2017). Programmable transcriptional repression in mycobacteria using an orthogonal CRISPR interference platform. Nat Microbiol 2, 16274.

Schneller, J.L., Lee, C.M., Bao, G., and Venditti, C.P. (2017). Genome editing for inborn errors of metabolism: advancing towards the clinic. BMC Med 15, 43.

She, P., Shiota, M., Shelton, K.D., Chalkley, R., Postic, C., and Magnuson, M.A. (2000). Phosphoenolpyruvate carboxykinase is necessary for the integration of hepatic energy metabolism. Mol Cell Biol 20, 6508–6517.

Shmakov, S., Smargon, A., Scott, D., Cox, D., Pyzocha, N., Yan, W., Abudayyeh, O.O., Gootenberg, J.S., Makarova, K.S., Wolf, Y.I., et al. (2017). Diversity and evolution of class 2 CRISPR-Cas systems. Nat Rev Microbiol 15, 169–182.

Sievers, F., Wilm, A., Dineen, D., Gibson, T.J., Karplus, K., Li, W., Lopez, R., McWilliam, H., Remmert, M., Soding, J., et al. (2011). Fast, scalable generation of high-quality protein multiple sequence alignments using Clustal Omega. Mol Syst Biol 7, 539.

Steinert, J., Schiml, S., Fauser, F., and Puchta, H. (2015). Highly efficient heritable plant genome engineering using Cas9 orthologues from Streptococcus thermophilus and Staphylococcus aureus. Plant J 84, 1295–1305.

Tsai, S.Q., Zheng, Z., Nguyen, N.T., Liebers, M., Topkar, V.V., Thapar, V., Wyvekens, N., Khayter, C., Iafrate, A.J., Le, L.P., et al. (2015). GUIDE-seq enables genome-wide profiling of off-target cleavage by CRISPR-Cas nucleases. Nat Biotechnol 33, 187–197.

Tycko, J., Barrera, L.A., Huston, N.C., Friedland, A.E., Wu, X., Gootenberg, J.S., Abudayyeh, O.O., Myer, V.E., Wilson, C.J., and Hsu, P.D. (2018). Pairwise library screen systematically interrogates Staphylococcus aureus Cas9 specificity in human cells. Nat Commun 9, 2962.

Yang, J., Kalhan, S.C., and Hanson, R.W. (2009). What is the metabolic role of phosphoenolpyruvate carboxykinase? J Biol Chem 284, 27025–27029.

Yang, Y., Wang, L., Bell, P., McMenamin, D., He, Z., White, J., Yu, H., Xu, C., Morizono, H., Musunuru, K., et al. (2016). A dual AAV system enables the Cas9-mediated correction of a metabolic liver disease in newborn mice. Nat Biotechnol 34, 334–338.

Yardeni, T., Eckhaus, M., Morris, H.D., Huizing, M., and Hoogstraten-Miller, S. (2011). Retro-orbital injections in mice. Lab Anim (NY) 40, 155–160.

Zetsche, B., Gootenberg, J.S., Abudayyeh, O.O., Slaymaker, I.M., Makarova, K.S., Essletzbichler, P., Volz, S.E., Joung, J., van der Oost, J., Regev, A., et al. (2015). Cpf1 is a single RNA-guided endonuclease of a class 2 CRISPR-Cas system. Cell 163, 759–771.

